# STIG: Generation and simulated sequencing of synthetic T cell receptor repertoires

**DOI:** 10.1101/2020.02.28.969469

**Authors:** Mark G. Woodcock, Dante S. Bortone, Benjamin G. Vincent

**Author notes:** UNC Division of Hematology/Oncology, Department of Medicine, Physician’s Office Building, 3rd Floor, 170 Manning Drive, CB #7305, Chapel Hill, NC, 27599, USA.

## Abstract

T cell receptor repertoire inference from DNA and RNA sequencing experiments is frequently performed to characterize host immune responses to disease states. Existing tools for repertoire inference have been compared across publicly available biological datasets or unpublished simulated sequencing data. Evaluation and comparison of these tools is challenging without common data sets created from a known repertoire with well-defined biological and sequencing characteristics. Here we introduce STIG, a tool to create simulated T cell receptor sequencing data from a customizable virtual T cell repertoire, with clear attribution of individual reads back to locations within their respective T-cell receptor clonotypes. STIG allows for robust performance evaluation of T cell repertoire inference and downstream analysis methods. STIG is implemented in Python 3 and is freely available for download at https://github.com/Benjamin-Vincent-Lab/stig

**Author summary:** As part of the acquired immune system, T cells are integral in the host response to microbes, tumors and autoimmune disease. These cells each have a semi-unique T cell receptor that serves to bind a set of antigens that will in turn stimulate that cell to perform its particular pro- (or anti) inflammatory role. This receptor is the product of DNA rearrangement of germline gene segments, similar to B cell receptor loci rearrangement, which provides a wide variety of potential T cell receptors to respond to antigens. At the site of an immune reaction, T cells can increase their number through clonal expansion and methods have been developed to analyze bulk genetic sequencing data to infer the individual receptors and the relative size of their clonal subpopulations present within a sample. To date, these methods and tools have been tested and compared using either biological samples (where the true quantitiy and types of T cells is unknown) or unshared synthetic datasets. In this paper I describe a new tool to generate biologically-inspired T-cell repertoires in-silico and generate simulated sequencing data from them.

## Introduction

Characterization of T cell receptor repertoires using next generation sequencing has allowed the adaptive immune system in various states of health and to be studied in fine detail. The number and clonal diversity of T cells can be measured to give insight to the host’s resting immune state and response to disease, as well as being potentially prognostic or predictive of response to therapy [1–7]. In bulk sequencing approaches, the sequences of T cell receptor chains must be inferred from read fragments. This process involves some uncertainty as individual reads may only partially fall into the receptor region of interest and read depth can limit detection of rare receptor clonotypes [8]. A number of tools have been developed to infer the underlying repertoire of T cell receptor chains from bulk sequencing data [8–12]. To properly evaluate these software tools and their underlying methodologies, a known “ground truth” repertoire must be constructed from which inferences can be compared. Existing tools for simulating TCR gene recombination lack features to mimic next-generation sequencing output (e.g. FASTQ-formatted output, DNA or RNA library preparation approaches, quality limitations) that might otherwise allow their direct use with inference tools.

There remains a need for tools that can create simulated TCR sequencing data from an underlying set of virtual T cell receptors. To that end, we developed Synthetic TCR Informatics Generator (STIG) to allow the creation of simulated sequencing data that reflects underlying T cell receptor biology, commonly used sequencing library approaches, and sequencing quality limitations. STIG is capable of generating sets of virtual T cell receptors (repertoires) with a customizable and biologically-inspired recombination model; created repertoires are used by STIG to generate simulated sequencing data reflecting multiple library preparation methods and user-defined fragment lengths and quality limitations. FASTQ-formatted outputs from STIG can be directly integrated with established T cell repertoire inference tools, and individual reads are labeled with the individual clonotype, receptor chain and relative location within the chain gene to allow easy quantification of data characteristics (e.g. read depth, CDR3 coverage per clone).

### Design and Implementation

STIG attempts to mimic T cell biology as closely as possible, internally modeling individual T cells with each possessing a TCR composed of two paired receptor chains (either alpha & beta, or gamma & delta). Clonal T cells with identical receptors are grouped together as a clonotype, and the set of all clonotypes is referred to as the repertoire. Creation of a simulated repertoire within STIG (outlined in Fig 1) begins with instantiation of a number of virtual clonotypes. Because the number of clonotypes and distribution of their sizes may vary by tissue sampled and the state of an individuals health or disease [13, 14], STIG supports multiple distributions to assign virtual T cells between clonotypes: even, pseudo-normal and skewed (e.g. logistic, see S1 Fig). Each clonotype is assigned a TCR receptor type (i.e. alpha/beta or gamma/delta) based on a ratio provided by the user before undergoing simulated V(D)J recombination. The recombination process is highly configurable, with the ability to define specific V segment odds and conditional probabilities for gene segments based on choice of prior segments (thus reflecting the nonlinear probabilities seen in biological TCR gene segment distributions [15]). Segment probabilities are customizable for every gene segment in all TCR chain types. Nucleotide addition and chewback during recombination can be modified from reasonable defaults. As transparency and flexibility of the recombination model are core to STIG’s utility, all recombination probabilities are exposed and adjustable through a well documented and user-editable configuration file. Users may provide custom segment allele sequences in FASTA-formatted files. The recombination results are tested to ensure results are in-frame, without early stops, and contain an appropriate CDR3 amino-acid motif, before being added to the repertoire: chain validation is crucial for simulating RNA-seq experiments, as the overwhelming majority of TCR chain RNA expression consists of chains with functional CDR3s [16, 17]. Options exist to enforce uniqueness between clonotypes at the level of the combined TCR chain pair, individual TCR chain, or individual CDR3. Generated repertoires are stored in a datafile alongside a comma-separated value (CSV) file containing statistical information about the repertoire, including the number clonotypes, quantity of cells in each clonotype, V and J region alleles utilized, and the DNA, RNA and CDR3 sequences of each TCR chain.

**Fig 1.**
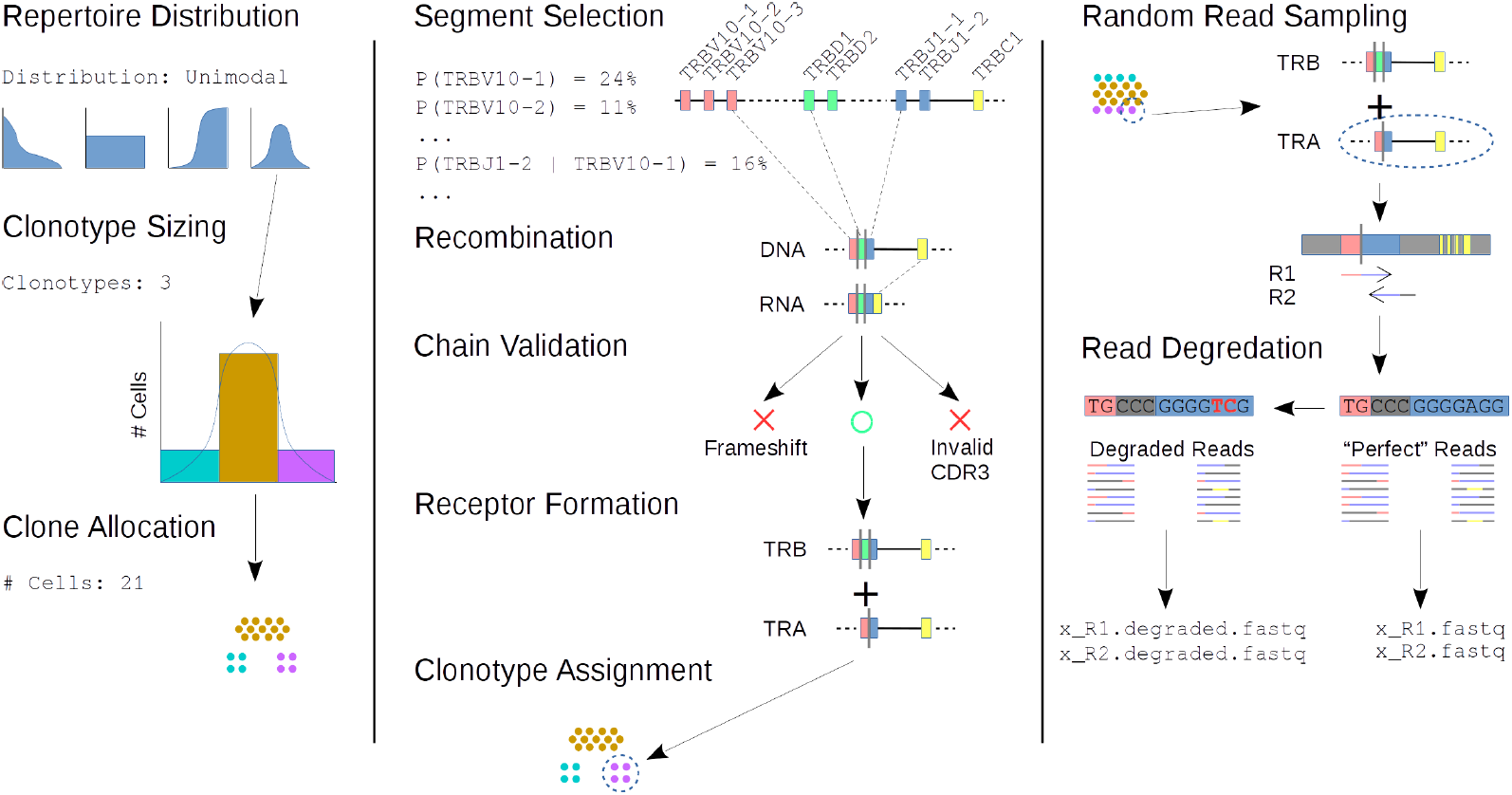
Graphical representation of repertoire generation and sequencing operations in STIG. Example parameters are listed in fixed-width font.

Created repertoires are subsequently used within STIG to generate simulated sequencing output. At its most basic, users specify the simulated sequencing approach (single end, paired end, or amplicon), nucleotide space (DNA or RNA-seq), and the number of reads to output. Users may further configure read and insert fragment lengths, with fixed or variable lengths defined by a customizable normal distribution (e.g. paired-end reads with a read length mean of 48, standard deviation from the mean of 8 nucleotides, and restrict read lengths to only those within 4 standard deviations from the mean.) Insert lengths are similarly flexible. Each read is randomly sampled by clone, TCR chain, and position within the chain gene; reads that fall partially outside the gene are completed using data from reference chromosome files to simulate the 5’ and 3’ untranslated regions. This sampling approach allows STIG to generate all possible reads that might include a portion of the TCR gene locus. STIG supports custom amplicon anchors and will attempt to align these in real-time during read generation. As the repertoire is sampled, perfect-quality reads are written to a FASTQ output with a comment line including the sampled clonotype ID, receptor chain, and relative position of the read within that chain. These machine-readable comment values allow users to associate individual reads back to the underlying TCR RNA or DNA and calculate parameters such as read depth and CDR3 coverage.

To simulate non-optimal read quality, STIG will optionally degrade the quality and nucleotide strings of generated reads. Several methods are provided: A fixed PHRED+33 string applied to all generated reads, a length-dependent logistic function, or a file-based approach where STIG duplicates quality strings from a user-provided FASTQ file. Read sequences are degraded based on the error probabilities implied by the paired PHRED+33 quality string before being written to a second set of output FASTQ files. The degraded reads are numbered and written in a one-to-one fashion, such that each degraded read can be linked back to a corresponding perfect quality read, clonotype and TCR location.

STIG is written in Python 3 and has two widely-adpoted Python packages as external requirements: pyyaml for parsing STIG’s YAML-formatted configuration file, and numpy for generating user-requested clone distributions. Correctness of the software’s operation is ensured by a set of unit tests as well as redundant internal self-consistency checks during runtime. STIG’s output has been tested using the published TCR inference tools MiXCR and TRUST to validate that the underlying simulated repertoire can be successfully reconstructed across STIG’s DNA, RNA and amplicon simulation modes; MiXCR’s reconstruction results in a simulated RNA-seq experiment are outlined below. Public releases are hosted on the project’s Github page (see “Availability and Future Directions” below) where bug reports and patches may be submitted.

### Comparison with existing tools

There are existing tools for TCR chain analysis and simulation that have some overlap with STIG’s functionality. IgoR [18] is a tool for inferring T cell alpha and beta chain (as well as Ig-heavy) V(D)J recombination models from sequencing data. It has the ability to generate DNA sequences from individual TCR chains based on an inferred (or user-provided) recombination model and allele data. Similarly, repgenHMM [19] was designed to infer an alpha or beta-chain recombination model from sequencing data using an internal hidden Markov model, and can generate synthetic DNA sequences for those chains with the likelihood of generation implied by the model. OLGA [20] was created to estimate probabilities for creation of user-provided alpha and beta chains (as well as Ig-heavy chains) within a given recombination model, using a dynamic programming approach to search across possible recombination paths.

While STIG does support creating T cell receptor chains based on a recombination model, its core utility — creating virtual T cell repertoires from which to generate realistic DNA or RNA-seq data — differs from other TCR analysis software (S1 Table). STIG’s use of a biologically-inspired model for T cell repertoires and receptors, as well as its support for RNA-space reads and multiple sequencing library approaches, is novel functionality compared to existing tools. By creating sequencing data from a set of T cell clones with a known recombination model, STIG could in fact be used to validate the inference operations of the above tools.

## Results

### MiXCR repertoire reconstruction

As a test use case, we measured the accuracy of the repertoire inference tool MiXCR using simulated RNA-seq data from STIG. We created three virtual repertoires with 150,000 cells between 250 clonotypes, each following a different distribution: Even (each clonotype with roughly equal numbers of cells), unimodal (many low and moderate population clonotypes, with a few very large clonotypes), and logistic (a large number of medium-sized clonotypes with thin tails of small or large clonotypes) (see S2 Fig). Each repertoire was used to generate successively larger numbers of FASTQ reads corresponding to the read-depths in Fig 2. MiXCR was run on the resulting outputs using the suggested RNA-seq settings (see S1 Text). To measure the “best case” accuracy we used only perfect quality reads from STIG. Uniqueness between the CDR3 portions of all chains within each repertoire was enforced using STIG’s repertoire creation options. The entire process of repertoire creation, read generation and MiXCR inference was repeated across 10 iterations to give the error bars shown in Fig 2.

**Fig 2.**
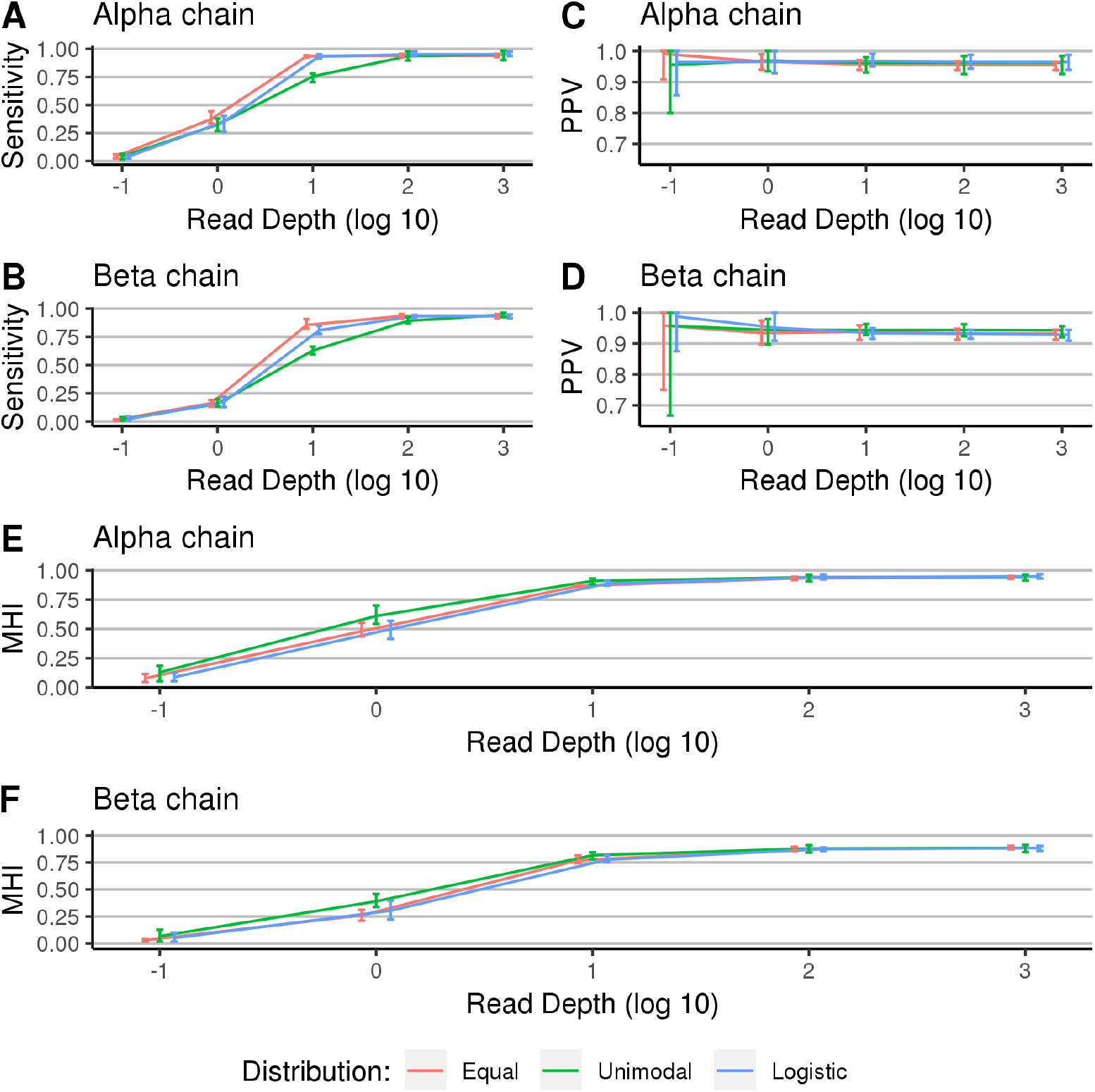
Accuracy of MiXCR reconstruction of three repertoires with different clonotype distribution. Error bars represent ranges across 10 iterations. Note the x-axis is log base 10 scale. (A-B) Sensitivity measured as the percentage of TCR chains from STIG’s underlying repertoire correctly inferred by MiXCR. (C-D) Positive predictive value measured as the percentage of TCR chains inferred by MiXCR that were contained in the underlying repertoire. (E-F) Horn-modified Morisita Overlap index comparing the similarities between the MiXCR inferred repertoire and the underlying repertoire. A value of one indicates full agreement between two distributions.

Our results show that MiXCR performs very favorably at read depths of 10 and above, and is sensitive to even low-population clonotypes at that read depth (Fig 2A,B). The sensitivity and positive-predictive value (PPV) indicate that MiXCR is able accurately discern the underlying CDR3 regions of both alpha and beta chains with narrow variation across the 10 iterations (Fig 2A-D). At sufficient read depths, MiXCR is able to accurately quantify the detected clonotypes (Fig 2E,F), based on the Horn-modified Morisita’s overlap index comparing the clonotype distributions between STIG’s output repertoire and MiXCR’s inferred repertoire. We noted the difference in the peak Morisita overlap index values between alpha and beta chains (Fig 2E,F), indicating that MiXCR was better at quantifying alpha chains over beta chains even at read depths exceeding that of typical RNA-seq experiments (i.e. 10^3^ coverage per base) [21]. These differing peak values were also observed when performing Kullback-Leibler divergence calculations between the inferred and underlying repertoires (see S2 Fig). Using data from the highest read depth analysis – chosen to minimize the effects of inadequate chain coverage – we compared the length of each chain against that chain’s contribution to the final Kullback-Leibler divergence score (Fig 3A,B). We hypothesize that any tendency to over-represent shorter CDR3s may be due to the higher probability of finding reads that unambiguously cover the CDR3 when that region is shorter. Further work is needed to confirm the relationship and any possible causality.

**Fig 3.**
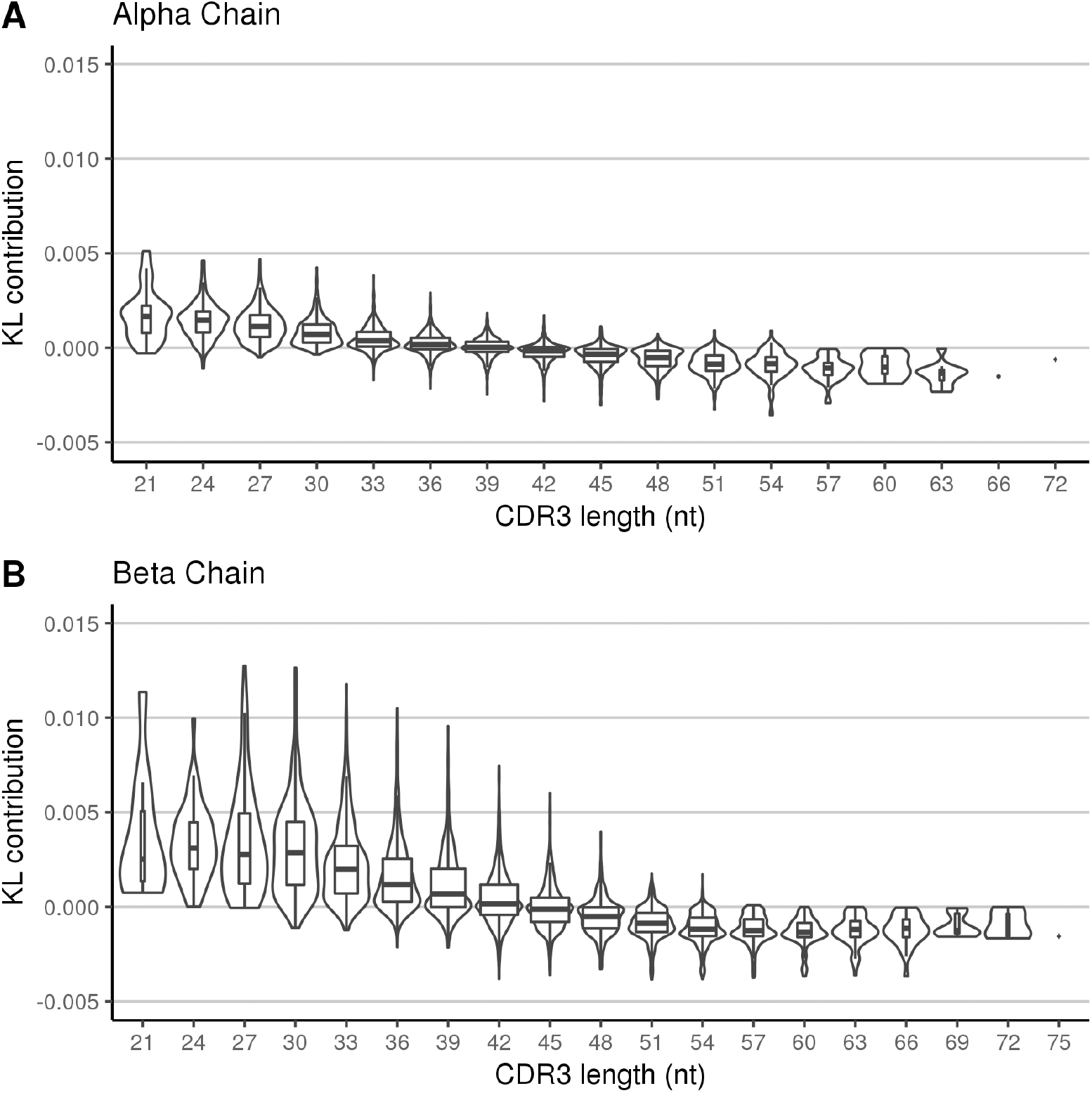
Contribution to the Kullback-Leibler divergence sum for each (A) alpha and (B) beta chain in the underlying STIG repertoire. Data represent inferences across all ten repetitions at the highest read depth analyzed (10^3^ coverage per base). Box widths are scaled to relative number of points at each length with whiskers covering values up to 1.5 times the interquartile range. Note that CDR3 lengths are not continuously distributed as they naturally must be a multiple of 3.

### Resource utilization

To measure CPU and memory requirements of repertoire generation, STIG was invoked to create repertoires of increasingly large numbers of clonotypes. Each invocation was repeated 10 times in total, with the maximum Resident-Shared Size (RSS) and CPU runtime recorded. Runtime and memory consumption increases linearly in proportion to the repertoire size (Fig 4), which is expected based on the design of STIG’s algorithm for generating T cell receptors. Each clonotype in the repertoire increases memory requirements by about 50Kb on average, as STIG stores the full DNA and RNA sequence of each TCR chain in memory. Storing this data in memory is a trade-off that allows for more efficient creation of simulated sequencing reads – STIG only accesses the on-disk chromosome reference files when reads fall into the 3’ or 5’ untranslated region.

**Fig 4.**
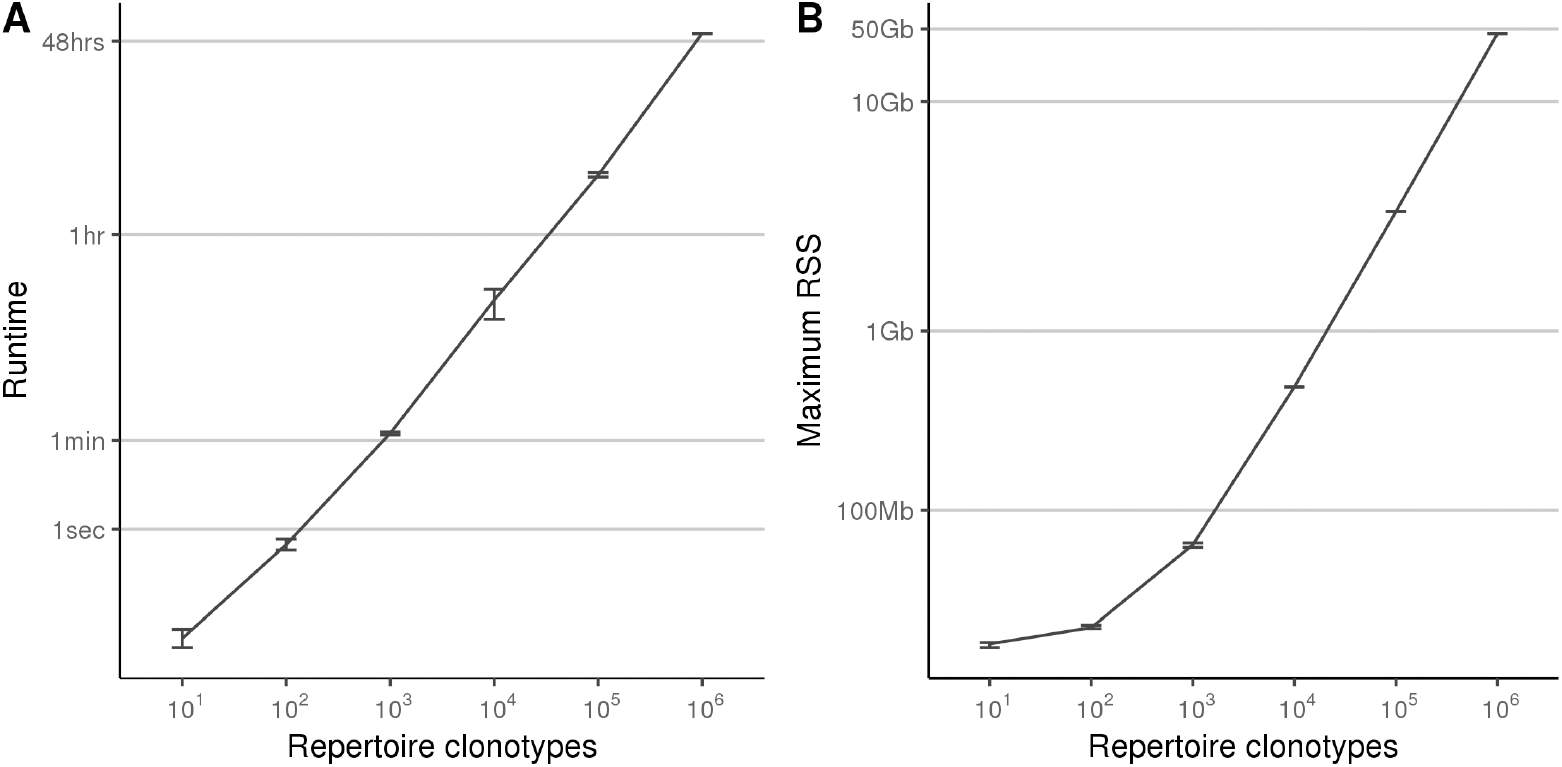
Resource utilization. (A) Average time to create a repertoire of simulated T cells, log scale. Error bars represent the maximum and minimum times across 10 repeated runs. Runtime is defined as the user + system time as reported by the ‘time’ command, as measured on an Intel Xenon ES-2680 2.4GHz CPU. (B) Average memory required to create a repertoire of simulated T cells, log scale. RSS is amount of memory allocated by STIG, as reported by the ‘ps’ command.

## Availability and future directions

STIG is written in Python 3 and available for download at the project repository (https://github.com/Benjamin-Vincent-Lab/stig). The functionality outlined in this manuscript is present in version 0.5.4. STIG is free software licensed under version 3 of the GNU General Public License.

STIG was created to simulate TCR sequencing data which mimics output from real-world sequencing approaches, and can be used directly with established TCR inference tools. The recombination model, repertoire characteristics and sequencing output parameters are adjustable through command line options, FASTA formatted allele files, and a text-based configuration file. STIG is species-agnostic and is distributed with a default recombination model based on human TCRs. Potential additions to the software include recombination models and alleles for other species. Applications of STIG include comparison of existing T cell repertoire inference tools, profiling approaches to estimate TCR chain pairings, and comparison of measures of T cell repertoire diversity and abundance.

## Acknowledgments

Computational work was performed in part using resources from the UNC Lineberger Bioinformatics Core.

## Supporting Information

**Supplementary Table 1.**
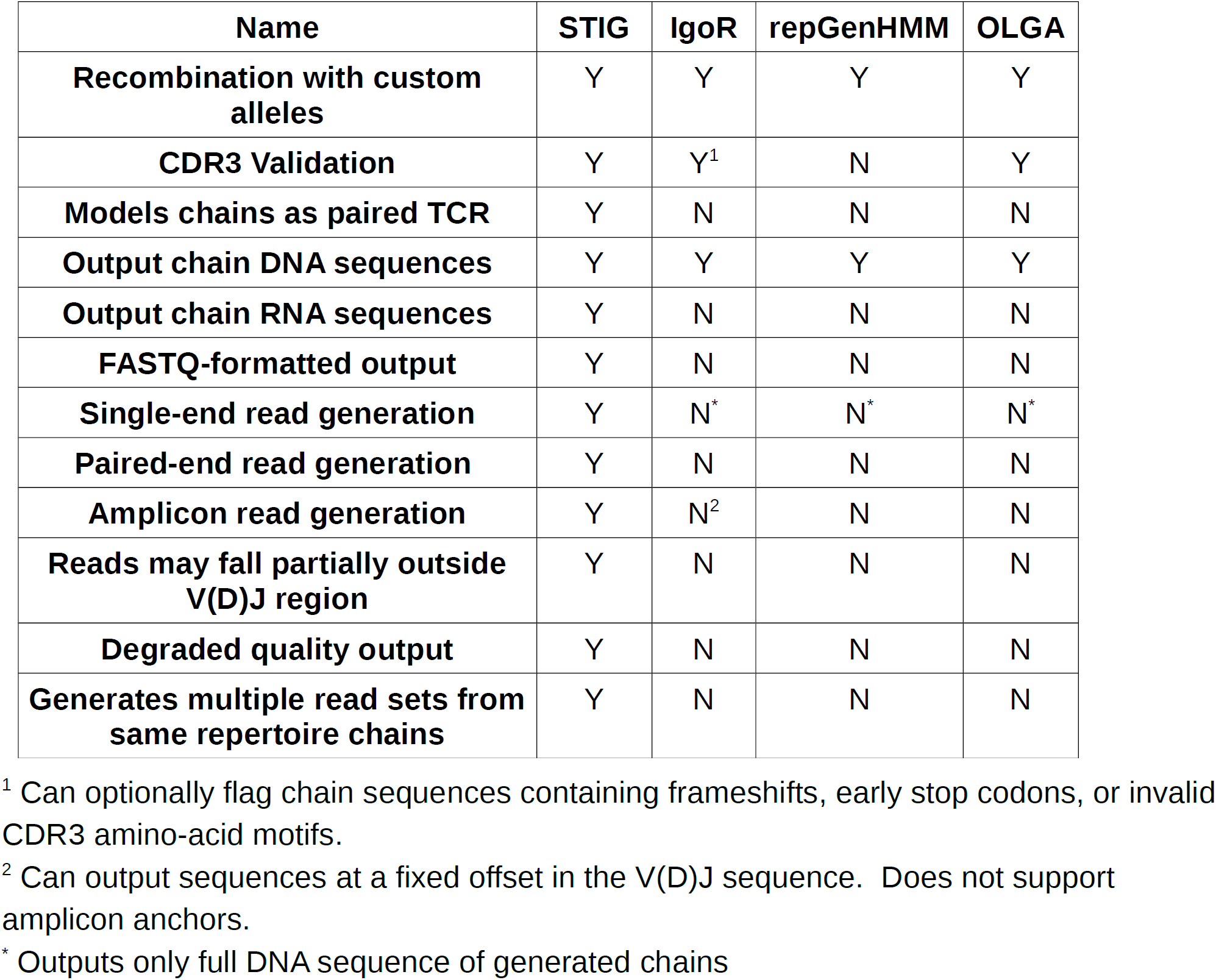
Feature Comparison of TCR Sequence Generators. Tools exist to output DNA sequences from recombination models but have limited support for simulating output seen in real-world sequencing approaches. “Models chains as paired TCR” refers to associating alpha-beta and gamma-delta chains to allow for realistic chain ratios during read generation.

### Supplementary Text 1

For analyzing paired-end RNA-seq data with MiXCR, we used the settings recommended for RNA-seq workflows by MiXCR’s authors (see manual at: https://mixcr.readthedocs.io/en/master/rnaseq.html#typical-analysis-workflow). The recommended settings for the alignment step as of this writing were:. The MiXCR version utilized was 2.1.9 (release date: 7 February, 2018).

### Supplementary Text 2

Effective read depths were calculated by trying to approximate a total RNA read depth of 0.1, 1, 10, 100 and 1000 reads per TCR region nucleotide (nt). The average alpha and beta chain gene RNA lengths were calculated from the generated repertoires yielding values of 816 and 933, respectively. Using 50nt paired reads with a 150nt insert length, that yields 816 + 933 + 600 nucleotides per chain in total (600 being the +150 nt * 2 per chain to account for reads that only partially fall into the 5’ or 3’ UTR regions), or 2349nt per virtual cell. With 100nt per read (50nt length, paired reads), one would need 23.49 reads per cell (or ∼11 per chain) for a read depth of 1. Thus, with 500 TCR chains between 250 clonotypes, our reads counts would be approximately 500 reads = 0.1 reads per base, 5000 reads = 1 read per base, and so on.

**Supplementary Figure 1.**
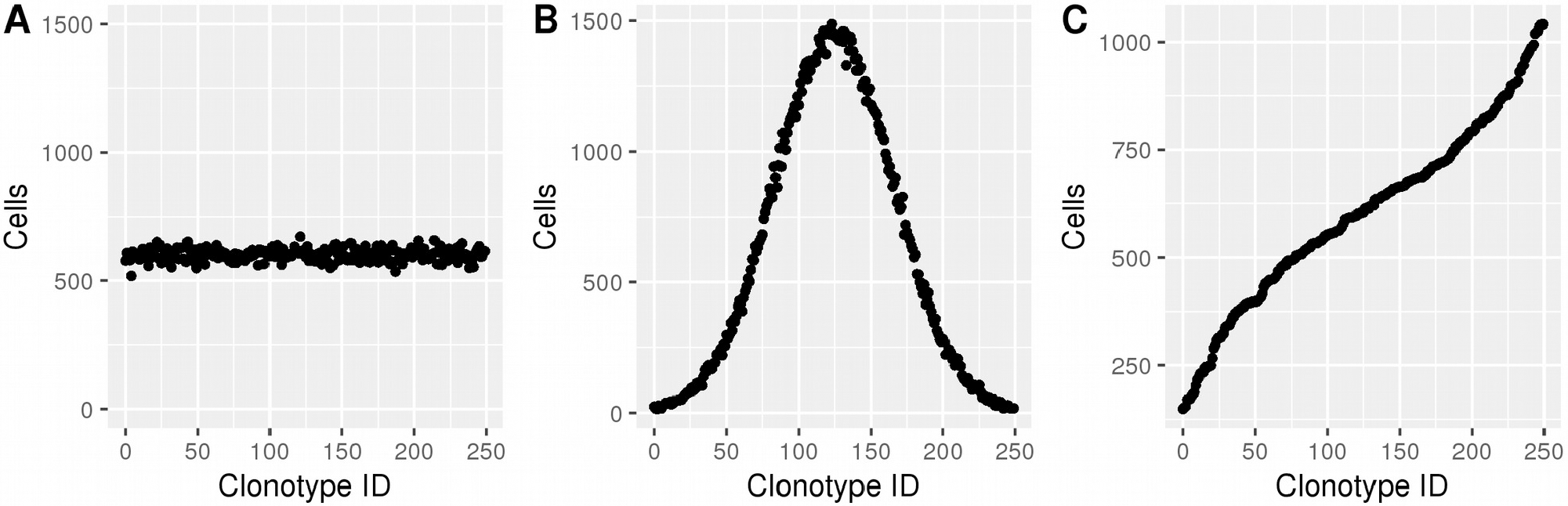
Example clonotype distributions in STIG. (A) The ‘equal’ option, in which cells are assigned to each clonotype with equal probability. (B) The ‘unimodal’ option. (C) The ‘logisticcdf’ option, which yields a truncated normal distribution when viewed as the number of clonotypes per population size.

**Supplementary Figure 2.**
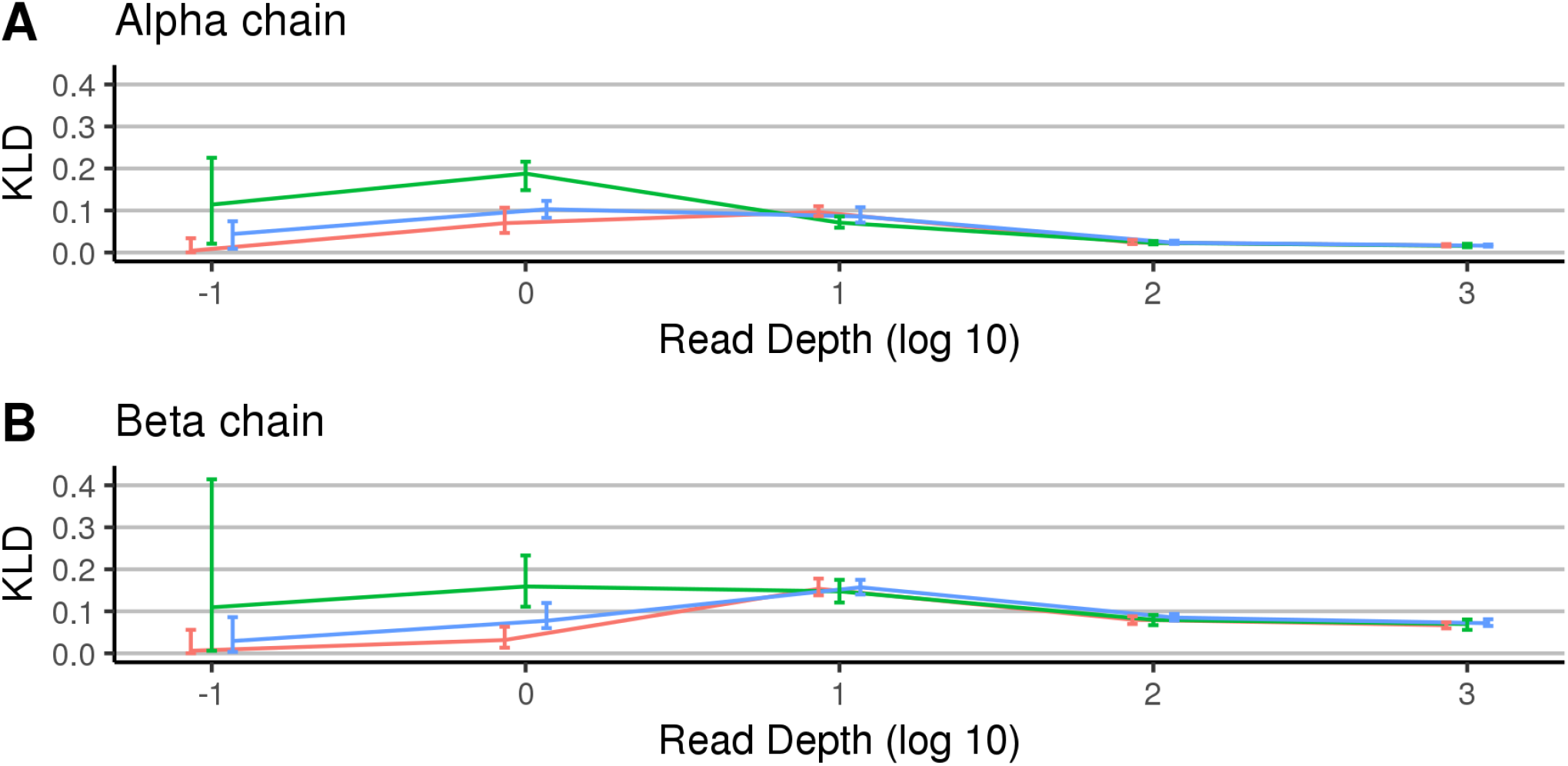
Kullback-Leibler divergence between STIG and MiXCR-inferred repertoires with different clonotype distributions. Error bars represent ranges across 10 iterations. Note the x-axis is log base 10 scale.

## References

1. Maini MK, Casorati G, Dellabona P, Wack A, Beverley PC. T-cell clonality in immune responses. Immunol Today. 1999;20(6):262–266.

2. Robins HS, Srivastava SK, Campregher PV, Turtle CJ, Andriesen J, Riddell SR, et al. Overlap and effective size of the human CD8+ T cell receptor repertoire. Sci Transl Med. 2010;2(47):47ra64.

3. Qi Q, Liu Y, Cheng Y, Glanville J, Zhang D, Lee JY, et al. Diversity and clonal selection in the human T-cell repertoire. Proc Natl Acad Sci USA. 2014;111(36):13139–13144.

4. Qi Q, Cavanagh MM, Le Saux S, NamKoong H, Kim C, Turgano E, et al. Diversification of the antigen-specific T cell receptor repertoire after varicella zoster vaccination. Sci Transl Med. 2016;8(332):332ra46.

5. Miller NJ, Church CD, Dong L, Crispin D, Fitzgibbon MP, Lachance K, et al. Tumor-Infiltrating Merkel Cell Polyomavirus-Specific T Cells Are Diverse and Associated with Improved Patient Survival. Cancer Immunol Res. 2017;5(2):137–147.

6. Ganesan AP, Clarke J, Wood O, Garrido-Martin EM, Chee SJ, Mellows T, et al. Tissue-resident memory features are linked to the magnitude of cytotoxic T cell responses in human lung cancer. Nat Immunol. 2017;18(8):940–950.

7. Jerby-Arnon L, Shah P, Cuoco MS, Rodman C, Su MJ, Melms JC, et al. A Cancer Cell Program Promotes T Cell Exclusion and Resistance to Checkpoint Blockade. Cell. 2018;175(4):984–997.

8. Warren RL, Nelson BH, Holt RA. Profiling model T-cell metagenomes with short reads. Bioinformatics. 2009;25(4):458–464.

9. Bolotin DA, Poslavsky S, Mitrophanov I, Shugay M, Mamedov IZ, Putintseva EV, et al. MiXCR: software for comprehensive adaptive immunity profiling. Nat Methods. 2015;12(5):380–381.

10. Kuchenbecker L, Nienen M, Hecht J, Neumann AU, Babel N, Reinert K, et al. IMSEQ–a fast and error aware approach to immunogenetic sequence analysis. Bioinformatics. 2015;31(18):2963–2971.

11. Gerritsen B, Pandit A, Andeweg AC, de Boer RJ. RTCR: a pipeline for complete and accurate recovery of T cell repertoires from high throughput sequencing data. Bioinformatics. 2016;32(20):3098–3106.

12. Li B, Li T, Pignon JC, Wang B, Wang J, Shukla SA, et al. Landscape of tumor-infiltrating T cell repertoire of human cancers. Nat Genet. 2016;48(7):725–732.

13. McHeyzer-Williams MG, Davis MM. Antigen-specific development of primary and memory T cells in vivo. Science. 1995;268(5207):106–111.

14. Laydon DJ, Bangham CR, Asquith B. Estimating T-cell repertoire diversity: limitations of classical estimators and a new approach. Philos Trans R Soc Lond, B, Biol Sci. 2015;370(1675).

15. Freeman JD, Warren RL, Webb JR, Nelson BH, Holt RA. Profiling the T-cell receptor beta-chain repertoire by massively parallel sequencing. Genome Res. 2009;19(10):1817–1824.

16. Redmond D, Poran A, Elemento O. Single-cell TCRseq: paired recovery of entire T-cell alpha and beta chain transcripts in T-cell receptors from single-cell RNAseq. Genome Med. 2016;8(1):80.

17. Singh M, Al-Eryani G, Carswell S, Ferguson JM, Blackburn J, Barton K, et al. High-throughput targeted long-read single cell sequencing reveals the clonal and transcriptional landscape of lymphocytes. Nat Commun. 2019;10(1):3120.

18. Marcou Q, Mora T, Walczak AM. High-throughput immune repertoire analysis with IGoR. Nat Commun. 2018;9(1):561.

19. Elhanati Y, Marcou Q, Mora T, Walczak AM. repgenHMM: a dynamic programming tool to infer the rules of immune receptor generation from sequence data. Bioinformatics. 2016;32(13):1943–1951.

20. Sethna Z, Elhanati Y, Callan J Curtis G, Walczak AM, Mora T. OLGA: fast computation of generation probabilities of B- and T-cell receptor amino acid sequences and motifs. Bioinformatics. 2019;doi: 10.1093/bioinformatics/btz035

21. Liu Y, Ferguson JF, Xue C, Silverman IM, Gregory B, Reilly MP, et al. Evaluating the impact of sequencing depth on transcriptome profiling in human adipose. PLoS ONE. 2013;8(6):e66883.

